# Identification of potential proteins and microRNAs in Multiple Sclerosis and Huntington’s diseases using in silico methods

**DOI:** 10.1101/2021.09.19.460949

**Authors:** Tarasankar Maiti, Souvik Chakraborty

## Abstract

Neurogenerative diseases like multiple sclerosis, Huntington’s disease are the major roadblocks in the way towards a healthy brain. Neurodegenerative diseases like multiple sclerosis and Huntington’s disease are affected by several factors such as environmental, immunological, genetics, and the worse scenario we can think of is that they are on the rise worldwide. Degenerative diseases specifically target a limited group of neurons at first resulting in the loss of specific functions associated with the specific part of the brain. The early diagnosis of these neurodegenerative diseases is important so that treatments can start from the early stages of these diseases. In this study, we have established a link between Multiple sclerosis and Huntington’s disease, and also we were able to establish the possible microRNAs that were connected to the expression of genes associated with these two diseases. In this present study, we analyzed the microarray datasets obtained from Gene Expression Omnibus and we identified 266 differentially expressed genes tried to identify using in silico methods the Hub genes involved in Multiple sclerosis and Huntington’s disease. After identifying the genes and proteins we tried to identify the microRNAs that are interacting with the Hub genes. In our study, we identified that the protein network has PTPRC, CXCL8, RBM25 proteins that have maximum connectivity. The top Hub genes are then subjected to a database that contains information concerning the microRNAs that are interacting with the Hub proteins as well as with each other. According to our study, the hsa-mir-155-5p has one of the highest degrees in the microRNA network. Our study will be useful in the future for the development of new drug targets for these neurodegenerative diseases.

## 1. Introduction

Neurodegenerative diseases (ND) like Huntington’s disease (HD) and Multiple Sclerosis (MS) are the common neuropathies. The prevalence of HD in the different studies is different. According to a study, it was estimated that 4.1 – 5.2 persons per 100,000 were living with the HD, but in another study, it was estimated that 2.4 – 8.4 persons per 100,000 were living with the disorder (Folstein et al., 1987; Kokmen et al., 1994; Reed & Chandler, 1958; Shokeir, 2008). HD is a rare autosomal dominant genetic neurodegenerative disorder that occurs due to CAG repeats in the Huntingtin gene (McCusker & Loy, 2017). CAG repeats in HD vary in a population and normally range from 11 to 33 but this number varies from 40 to 72 in HD patients (Zühlke et al., 1993). HD affects several areas in the brain that includes basal ganglia, hypothalamus, subthalamic nuclei, globus pallidus, substantia nigra, and also cerebellum (Eidelberg & Surmeier, 2011). Symptoms of Huntington’s disease include behavioral changes, cognitive impairment, and dysphagia, but the disease’s hallmark is uncontrollable spasmodic contractions known as Huntington’s chorea (Myers, 2004).

In 2016, 2.5 million people approximately were affected by MS but in 2020 this number has increased to 2.8 million approx (Dilokthornsakul et al., 2016; Walton et al., 2020). Typically the age of onset for the disease is 20 to 40 years but in almost 10 % of the cases, the symptoms start at an early age of 18 (Goldenberg, 2012). It has been seen in studies that MS is more prevalent in females than in males (Dilokthornsakul et al., 2016). MS is caused by an autoimmune reaction mediated by the CD4+ T cells. The involvement of the adaptive immune system leads to the formation of plaques in the CNS which are made of immune system cells, demyelinated axons, and an abnormal increase of astrocytes in the area (Gandhi et al., 2010). The microglial cells, the main antigen-presenting cells of the brain are activated when the blood-brain barrier is damaged due to the activation of T-cells. B cells also play a major role in the development of the disease and this is evident from the fact that B cell follicles in the meninges are highly active during the early onset of MS (Häusser-Kinzel & Weber, 2019).

Genetic control of both these disorders is an important player in the development and aggravation of these NDs. Despite a large number of studies, the treatment plans for both HD and MS remain symptomatic (Adam & Jankovic, 2008; Henze et al., 2006). Even though much study has been done to understand the underlying molecular pathways of NDs, there is still a lot more work to be done, because numerous brain areas are affected and due to this NDs must have common pathway convergence (Armstrong et al., 2005). Both HD and MS is increasing in an unprecedented way but there are no concrete ways to control them and therefore scientific studies like these should be conducted to unravel the mysteries of both these NDs. For the discovery of new and better therapeutic targets, a greater knowledge of these convergence pathways are critical. In this present study, we have analyzed the common pathways associated with HD and MS using publicly available microarray data (HD GSE 1767, MS GSE 21942) obtained from Gene Expression Omnibus (GEO) repository (Edgar, 2002). We have also studied the different microRNAs associated with the hub genes in the protein-protein network.

## 2. Materials and Methods

### 2.1. Dataset Selection

Datasets related to neurodegenerative diseases (NDs) are obtained from the Gene Expression Omnibus (GEO). The microarray gene expression data for Huntington’s disease (HD) is GSE1767 which used Affymetrix GeneChip U133A and Amersham Biosciences CodeLink chips contain data from blood (Borovecki et al., 2005). We have selected 12 samples from healthy individuals and 12 HD samples out of 12 symptomatic, 5 late presymptomatic patients. The dataset for Multiple Sclerosis (MS) is GSE21942 which used Affymetrix GeneChip Human Genome U133 Plus 2.0 Array containing data from peripheral blood mononuclear cells and we have selected 10 normal and 10 MS patients (Kemppinen et al., 2011).

### 2.2. Processing of Data and selection of Differentially Expressed Genes

The two datasets namely GSE1767 for HD and GSE21942 for MS are analyzed using the GEO2R tool (available at http://www.ncbi.nlm.nih.gov/geo/geo2r/) provided by GEO (Barrett et al., 2012). GEO2R creates a text file with a list of differentially expressed genes in disease vs. control format, using Bioconductor packages such as GEO query and limma R. LogFC values, p values, adjusted p values, gene symbols, gene IDs, and gene titles are included in this text file retrieved from GEO2R. The Differentially Expressed Genes (DEGs) having p-value greater than 0.05 were not considered for the study and the genes have a fold change value (logFC) greater than equal to 0.5 for both HD and MS.

### 2.3. Common DEG Identification and Network analysis

Fun Rich, a desktop-based tool was used to finding out the common DEGs in these two disorders (Pathan et al., 2015). For the development of a gene network, the online software called GeneMANIA was used (*GeneMANIA*, n.d.).The common DEGs were imported to STRING (Search Tool for the Retrieval of Interacting Genes) available at https://string-db.org/ and the protein-protein interaction (PPI) network was created using the information on the STRING database (Szklarczyk et al., 2019). The protein-protein network thus obtained from STRING was imported to Cytoscape directly from STRING (Shannon, 2003). The gene in the DEGs list with maximum connectivity to other genes (called hub genes) was found using a Cytoscape plugin called the CytoHubba (Chin et al., 2014). Enrichr an online software was then used for enrichment analysis (Chen et al., 2013). The DEGs which showed the maximum connection to other genes in the PPI were further considered microRNA interactions using online software called miRNet (Fan et al., 2016).

## 3. Results

In this present study, we used the microarray data available from Gene Expression Omnibus for two NDs, Multiple Sclerosis and Huntington’s disease. We studied the differentially expressed genes, nature of protein-protein interactions, identified the hub genes, and also identified the mi RNAs associated with the top 10 hub genes. The data used for this study has been normalized (Fig 1).

**Fig. 1:**
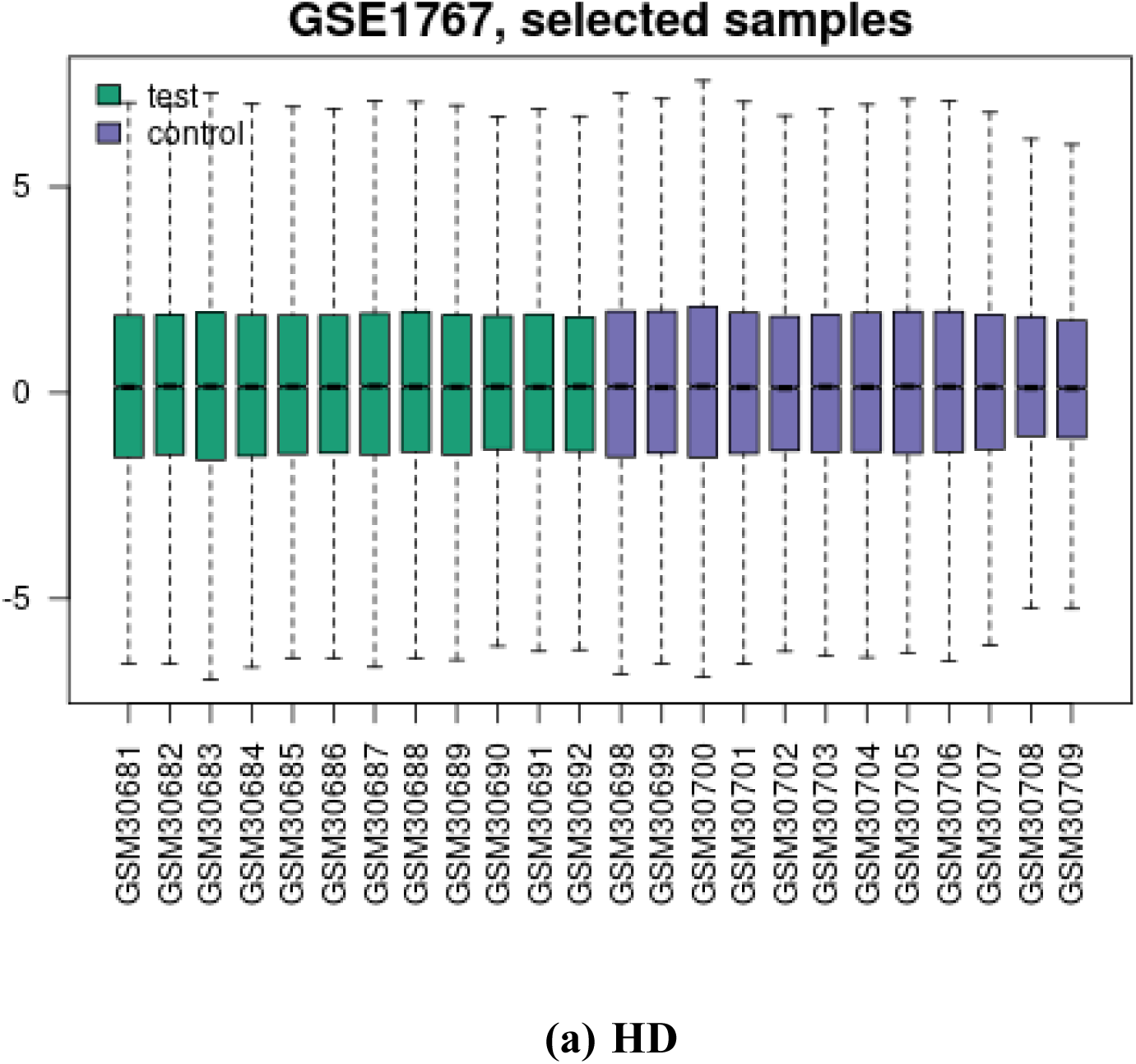
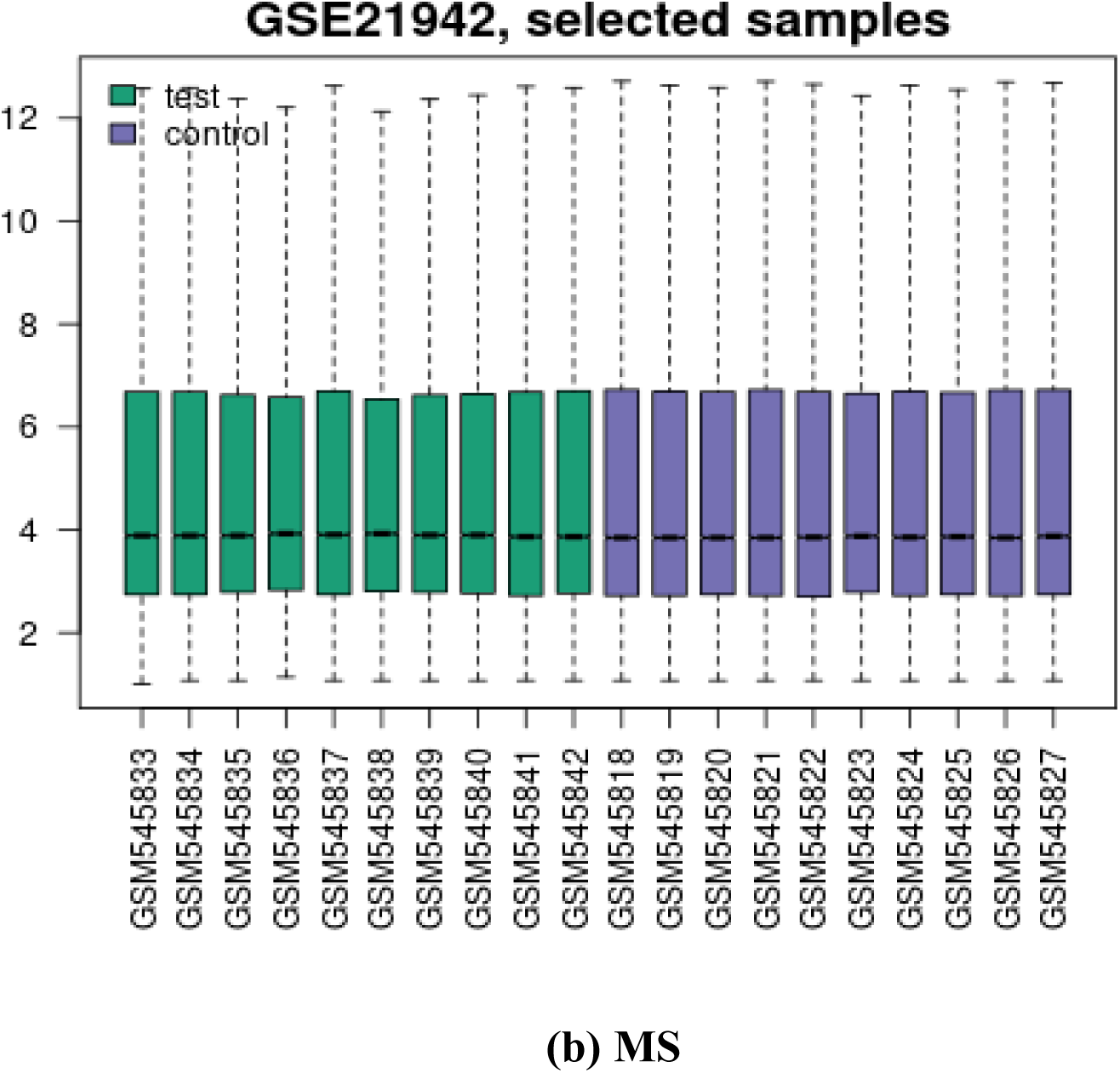
Box plots of HD and MS representing the normalization of data. The distribution of Control versus Diseased samples was depicted after data normalization with the GEO2R program. Each box plot represents the differential gene expression value for single patients.

**Fig. 2:**
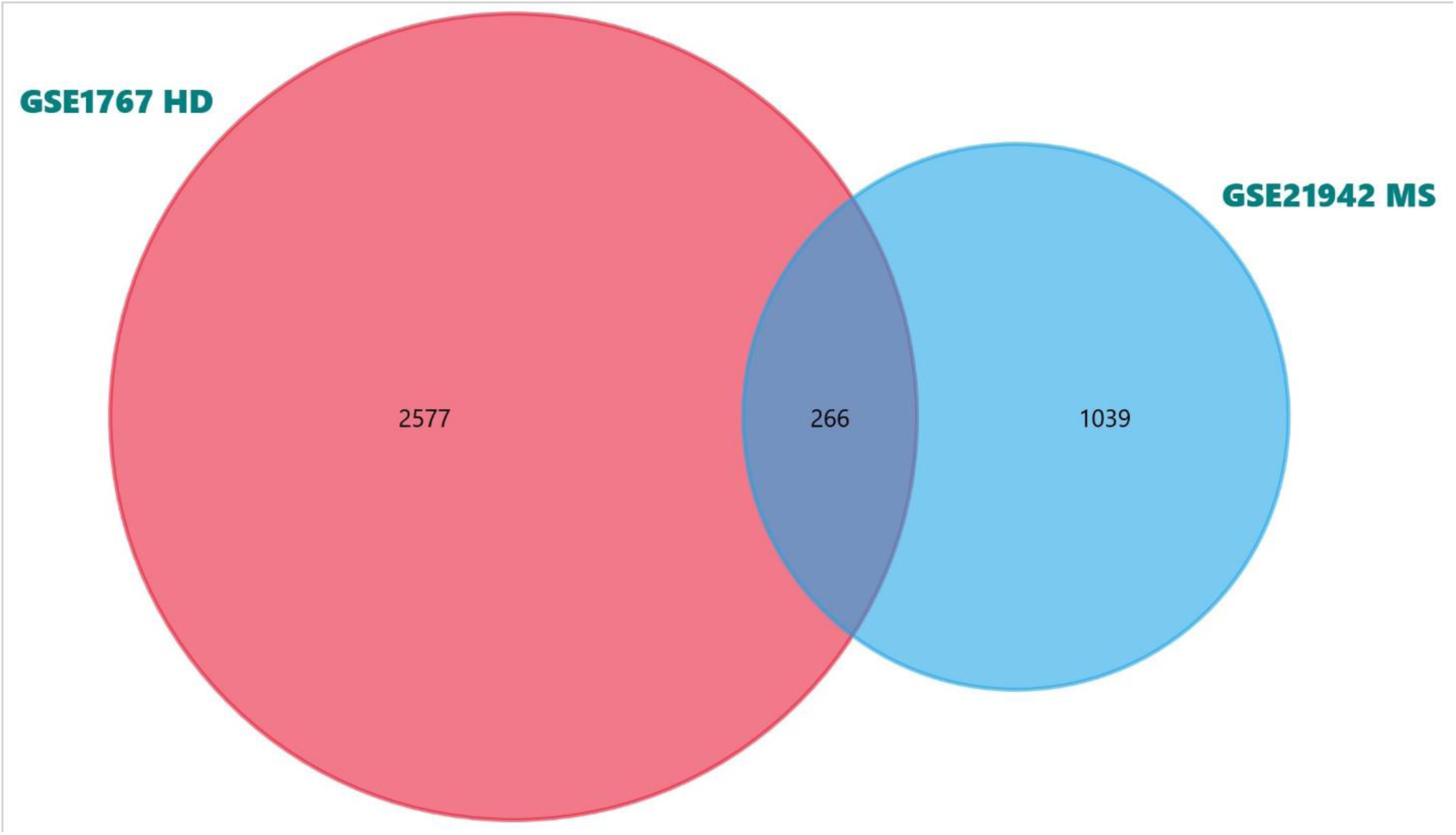
Venn diagram which represents the number of common differentially expressed genes in both the datasets, GSE21942 for MS and GSE1767 for HD.

### 3.1. Identification of Differentially expressed genes in two datasets

In the dataset GSE21942 out of 54675 genes, 1039 genes were found to be differentially expressed, and in the dataset GSE1767 out of 17932 genes, 2577 genes were differentially expressed which passed the set cut off of p-value 0.05 and logFC value of 0.5.

### 3.2. Finding the common DEGs in these datasets

The search for shared DEGs is necessary to determine whether the illness development processes in both of these disorders are identical. We observed that 266 genes were shared between the two datasets.

### 3.3. Interaction between common DEGs

The interaction among the common DEGs was visualized using the online version of GeneMANIA software (Fig 3). A dense gene network existed for the DEGs.

**Fig. 3:**
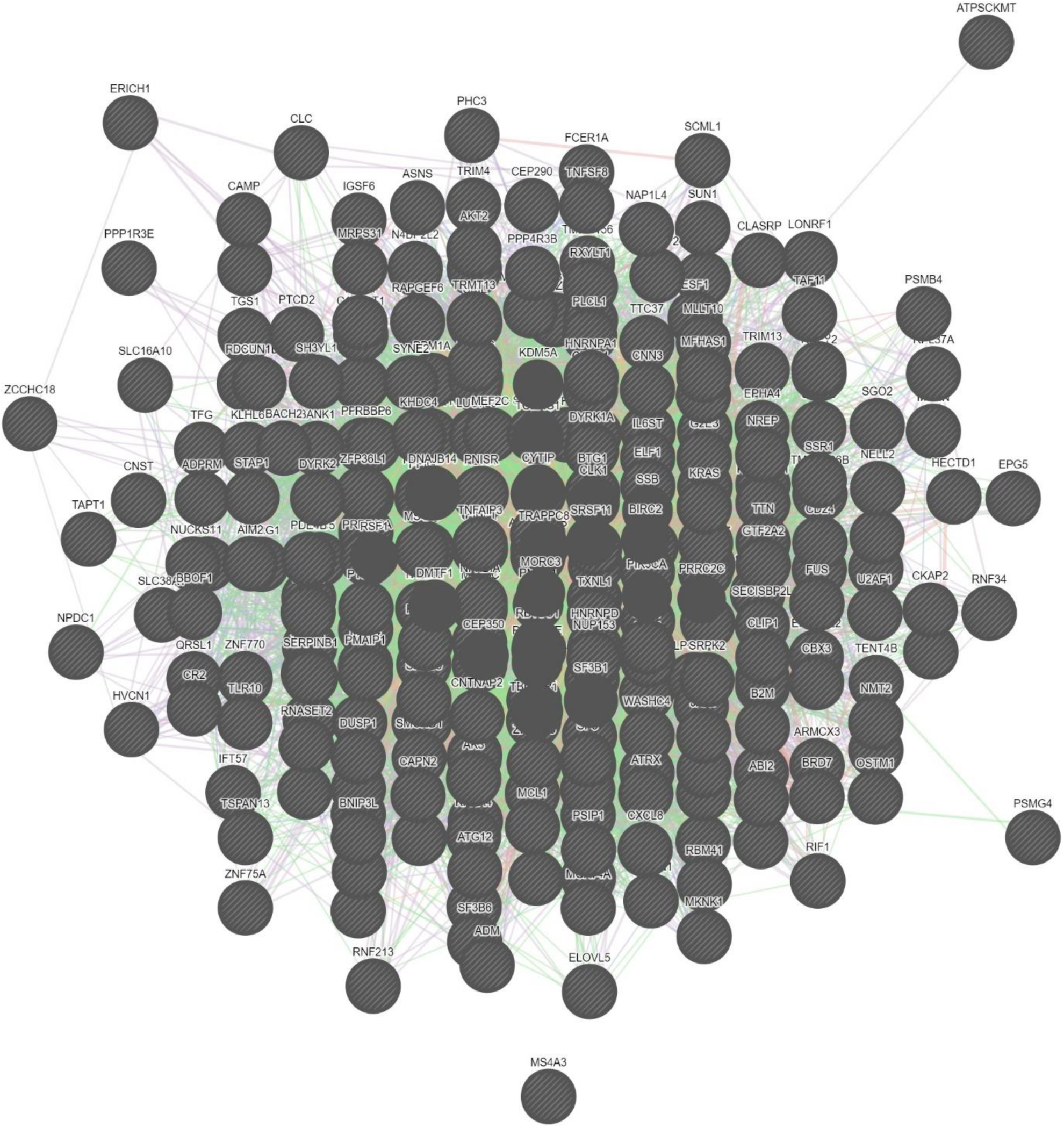
GeneMANIA network of all the differentially expressed genes (DEGs) in the datasets for HD and MS. Here all the genes which are common to both the datasets of HD and MS are represented.

The list of common DEGs is now subjected to the STRING database which uses Markov Clustering Algorithm for developing a protein-protein interaction (PPI) network (Fig 4).

**Fig. 4:**
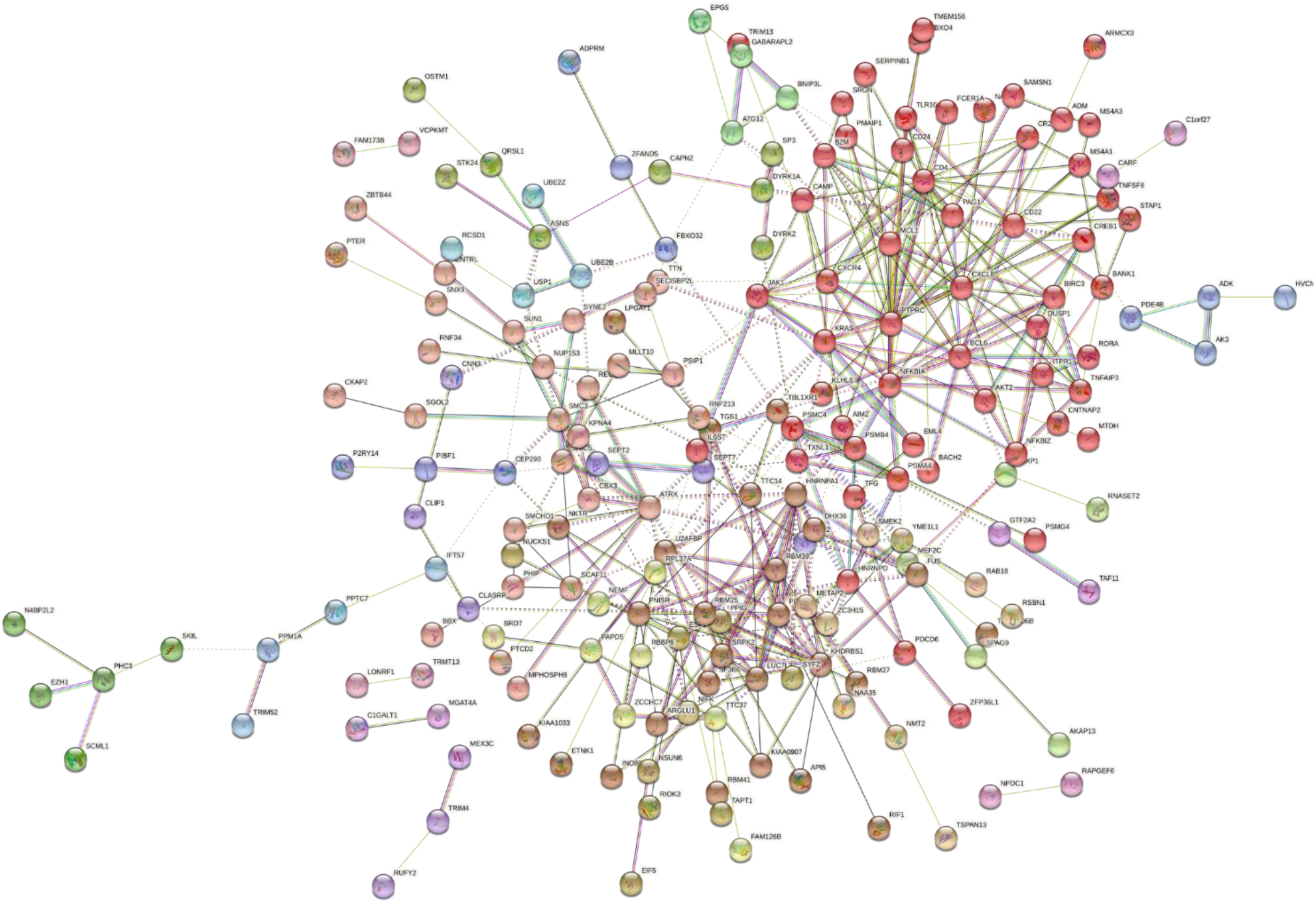
The protein-protein interaction network was developed by using the MCL clustering method. For this network, HD and MS DEGs were used. In this network, the interaction score was set at medium confidence (0.400).

The PPI had some proteins which do not interact at all while there are some interesting connections among many proteins in the network which had connections to other proteins as well. For identification of Hub genes, the PPI was imported to Cytoscape software and the CytoHubba plugin was used for visualizing and calculating the degree of centrality of all the hub genes (Table 1).

**Table-1:** Top ten Hub proteins in the PPI network obtained from the DEGs of the datasets for HD and MS with their degree of centrality.

| **Name of the protein** | **Degree score** |
| --- | --- |
| PTPRC | 19 |
| CXCL8 | 17 |
| RBM25 | 17 |
| KRAS | 16 |
| HNRNPA1 | 16 |
| NFKBIA | 15 |
| MCL1 | 14 |
| CXCR4 | 14 |
| BCL6 | 13 |
| KHDRBS1 | 13 |

The top ten hub genes were considered for constructing a network of PPI (Fig 5). It was seen that the Protein Tyrosine Phosphatase Receptor Type C (PTPRC) had the highest degree (Degree 19) followed by the Interleukin 8 (CXCL8) and RNA binding motif protein 25 (RBM25) with a degree of 17, Kirsten rat sarcoma viral oncogene homolog (KRAS) and Heterogeneous nuclear ribonucleoprotein A1 (HNRNPA1) had a degree of 16, NFKB inhibitor alpha (NFKBIA) with a degree of 15, MCL1 and CXC chemoreceptor type 4 (CXCR4) both had a degree of 14, BCL6 a transcription repressor, in the PPI and the protein KH RNA Binding Domain Containing, Signal Transduction Associated 1 (KHDRBS1) had the lowest degree (degree = 13). All the above-said genes were visualized properly when the interaction score in the STRING was set to high confidence (0.700).

**Fig. 5:**
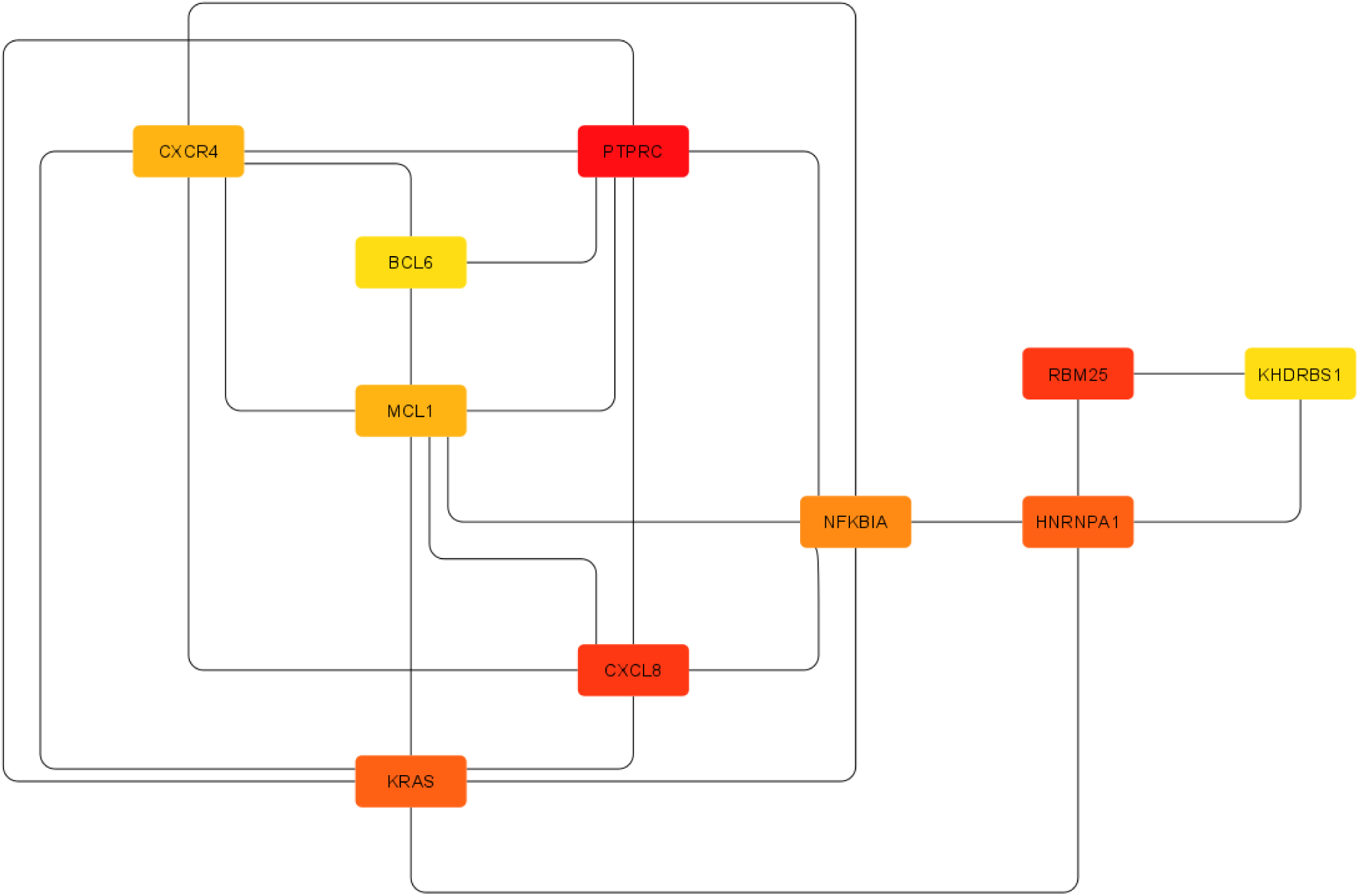
Network of top ten hub genes which was obtained using Cytoscape. The colour code represents the degree here, the red colour denotes the highest degree of centrality, orange denotes medium degree and yellow denotes the lowest degree of centrality.

PPI network showing the top ten hub genes was then obtained by setting the interaction score to high confidence (0.700) (Fig 6).

**Fig. 6:**
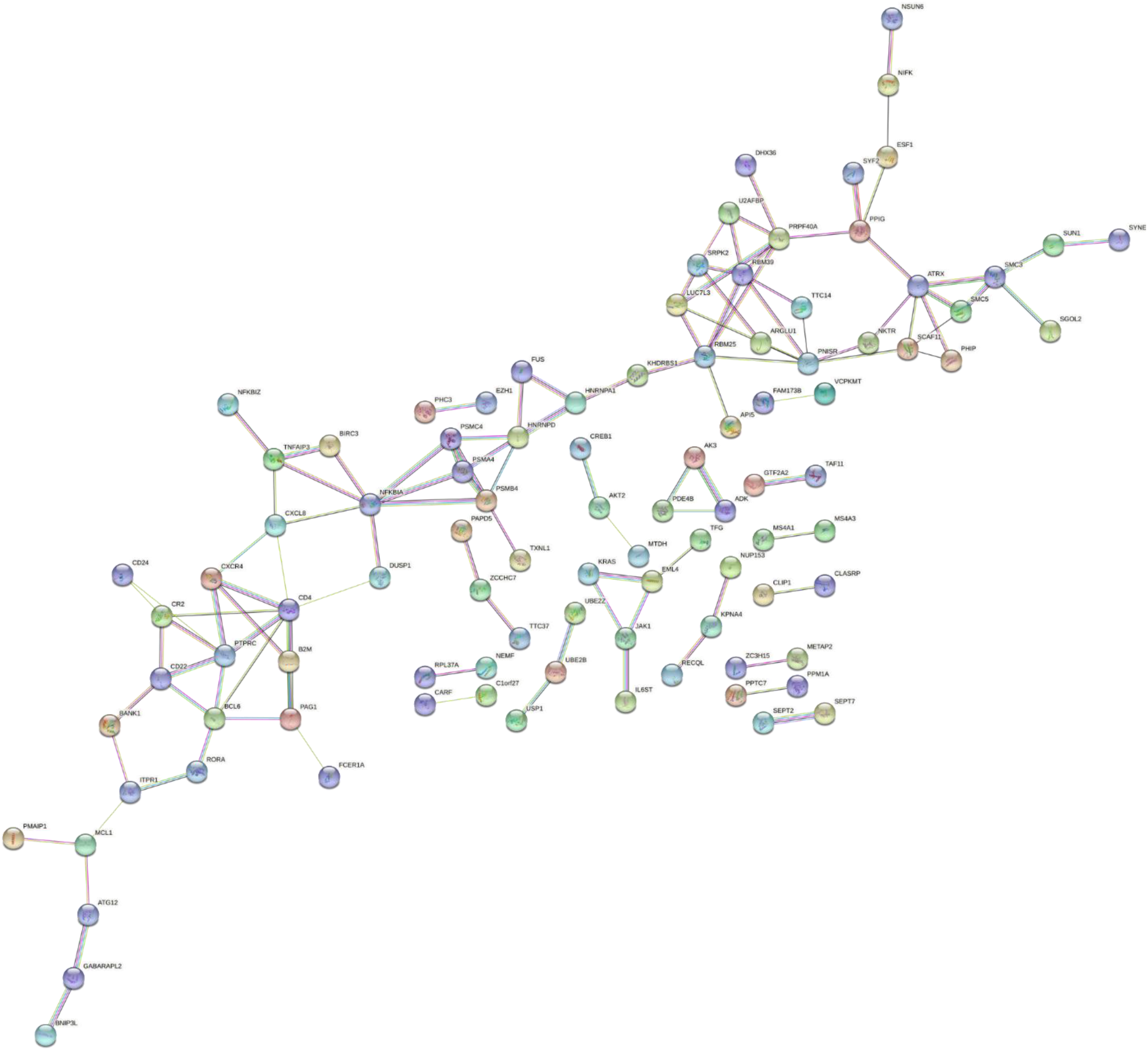
The PPI network showing the hub genes. This particular network was developed by setting the interaction score to high confidence (0.700), a high confidence score was used to reduce the interactions and to clearly state the Hub genes interacting with other genes.

### 3.4 Identification of miRNA and transcription factors associated with DEGs

The top ten hub genes were submitted to the miRNet, an online platform for finding out the interaction of genes with miRNAs. A network was generated which shows the interaction between DEGs,miRNAs, and transcription factors (Fig 7). 19 miRNAs and seven transcription factors were found to be associated with the 10 DEGs. In the network, the gene with maximum connectivity is MCL1 (degree 17) followed by the CXCL8 (degree 16). The miRNA called hsa-mir-155-5p has the highest connectivity in the network (degree 8) followed by miRNA hsa-mir-19a-3p (degree 5). The names of miRNA and the transcription factors associated with these ten DEGs are listed in Table 2.

**Table-2:** List of miRNAs and transcription factors together with 10 DEGs with their degree of centrality common to MS and HD.

| **Names of DEGs, miRNA, and transcription factors** | **Degree** |
| --- | --- |
| MCL1 | 17 |
| CXCL8 | 16 |
| NFKBIA | 10 |
| KRAS | 9 |
| hsa-mir-155-5p | 8 |
| KHDRBS1 | 8 |
| CXCR4 | 7 |
| BCL6 | 6 |
| hsa-mir-19a-3p | 5 |
| hsa-mir-93-5p | 5 |
| hsa-mir-96-3p | 5 |
| NFKB1 | 4 |
| RELA | 4 |
| hsa-mir-19b-3p | 4 |
| hsa-mir-25-3p | 4 |
| hsa-mir-181a-5p | 4 |
| HNRNPA1 | 4 |
| RBM25 | 4 |
| MYC | 3 |
| STAT3 | 3 |
| hsa-mir-17-5p | 3 |
| hsa-mir-20a-5p | 3 |
| hsa-mir-146a-5p | 3 |
| PTPRC | 3 |
| ATF4 | 2 |
| DDIT3 | 2 |
| ERG | 2 |
| hsa-mir-21-5p | 2 |
| hsa-mir-106b-5p | 2 |
| hsa-mir-18a-3p | 2 |
| hsa-mir-940 | 2 |
| hsa-mir-212-3p | 2 |
| hsa-mir-18a-5p | 2 |
| hsa-mir-126-5p | 2 |
| hsa-mir-20a-3p | 2 |
| hsa-mir-33a-3p | 2 |

**Fig. 7:**
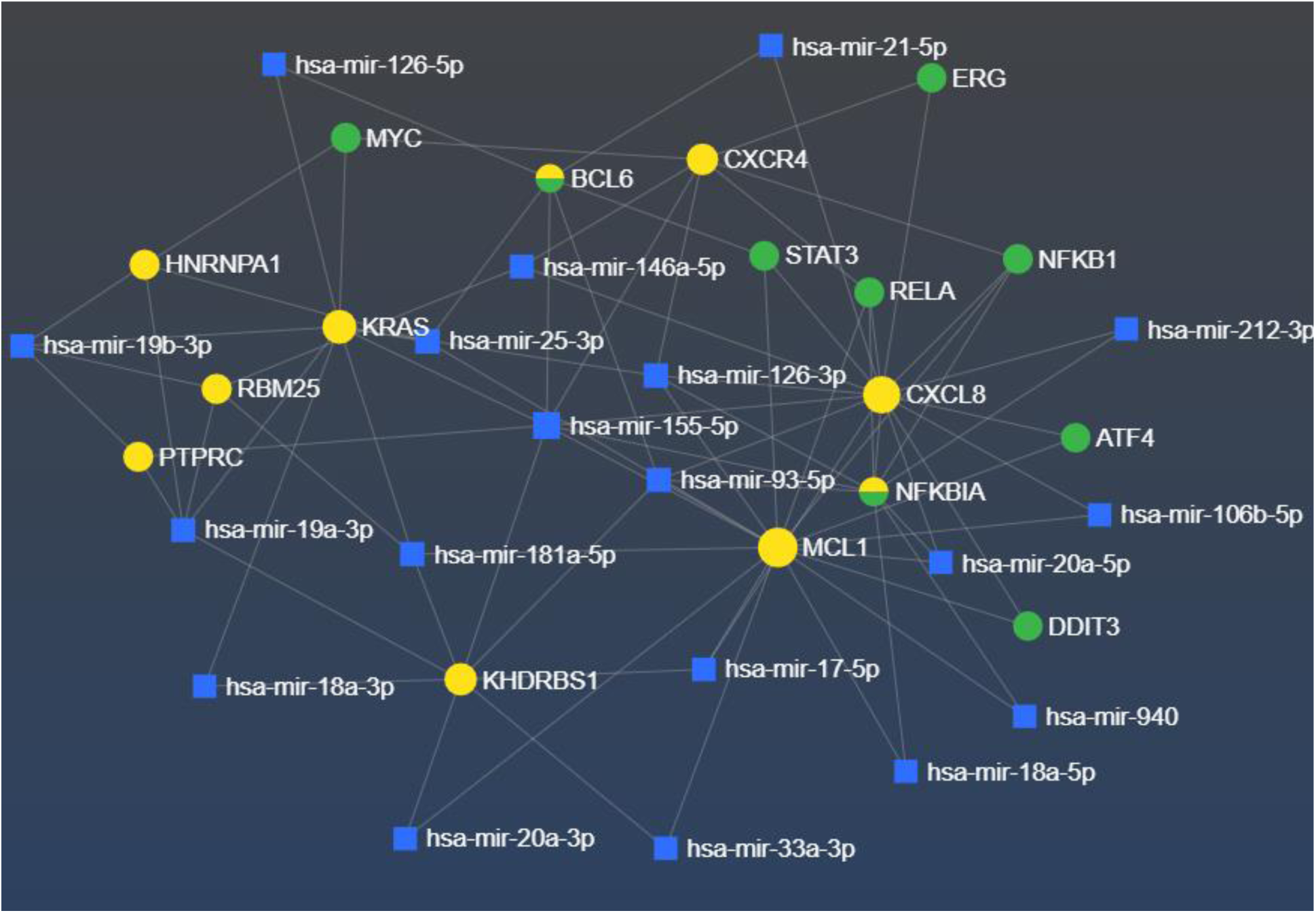
The network shows the interaction among DEGs, miRNAs, and transcription factors generated using miRNet. The genes are represented in yellow, the miRNA is represented in blue, transcription factors in green and transcription factors with identical gene and protein names are dual-colored (i.e. green and yellow-colored). The genes MCL1 and CXCL8 have maximum connectivity (degree 17 and 16 respectively) and the miRNA hsa-mir-155-5p and hsa-mir-19a-3p are the ones with maximum connectivity (degree 8 and 5).

### 3.5. Common GO processes and KEGG pathways for the DEGs associated with HD and MS

For the enrichment analysis, the DEGs were submitted to the online software, Enrichr, and GO biological processes, GO molecular function, GO cellular component and KEGG pathways options were used for the enrichment analysis.

#### 3.5.1. GO biological processes enriched which are common

The biological process that is significantly most enriched is the negative regulation of lymphocyte activation (GO:0051250) followed by the regulation of germinal center formation(GO:002634) and regulation of I kappa B kinase/NF kappa B signaling (GO:0043122). The graphical representation of the GO biological processes common to the set of DEGs submitted has been represented graphically (Fig 8a)

**Fig. 8:**
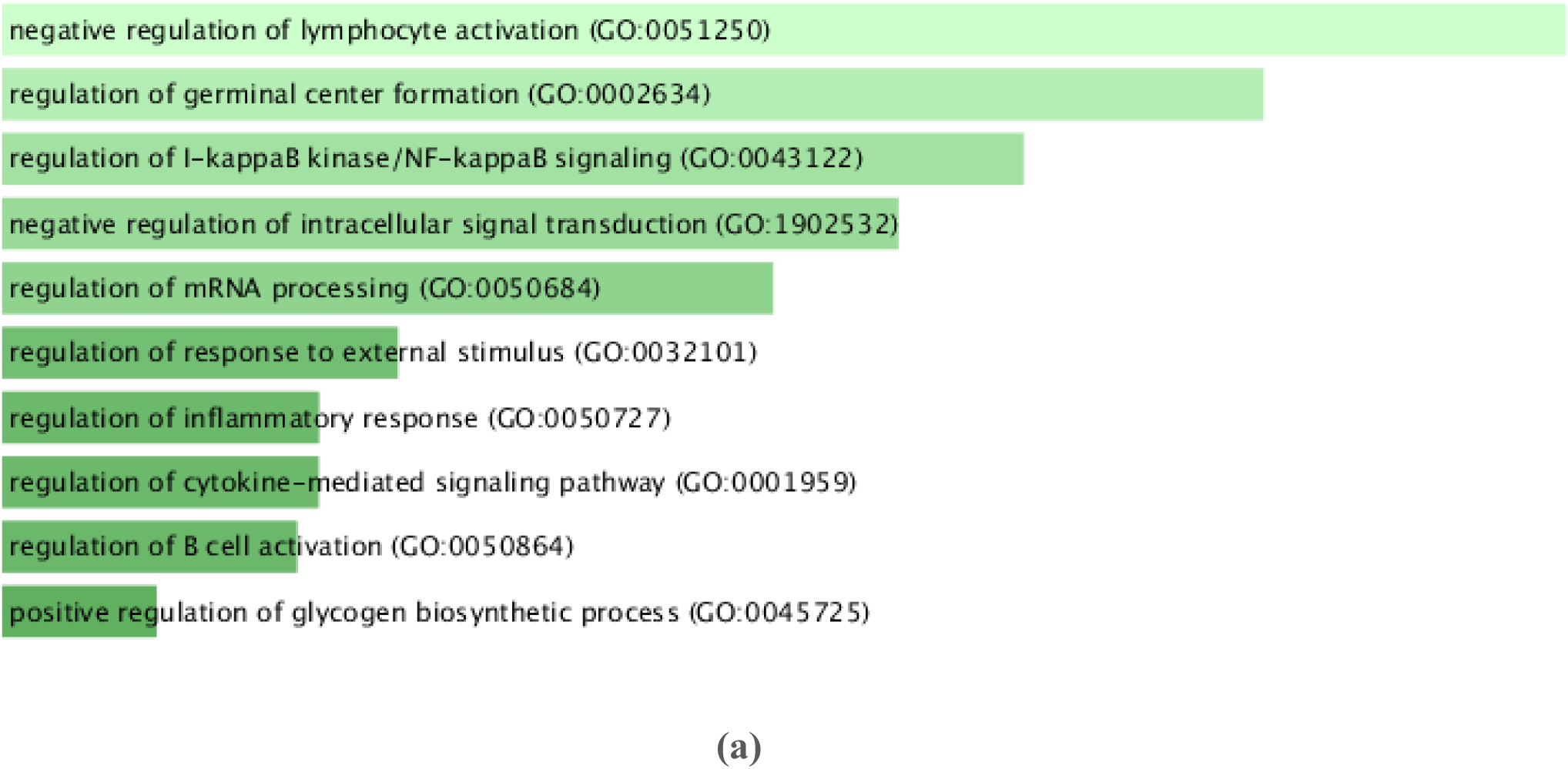
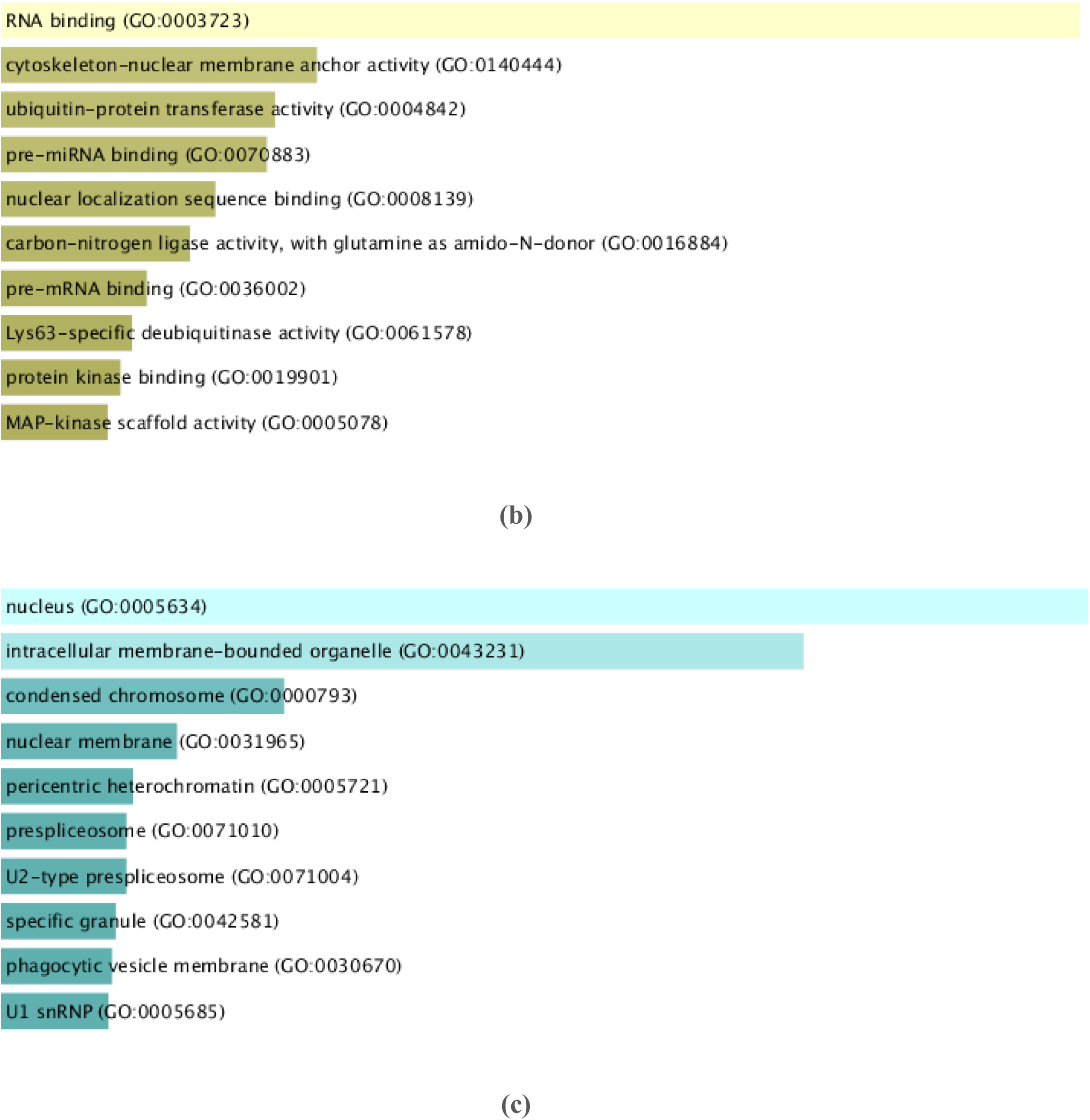
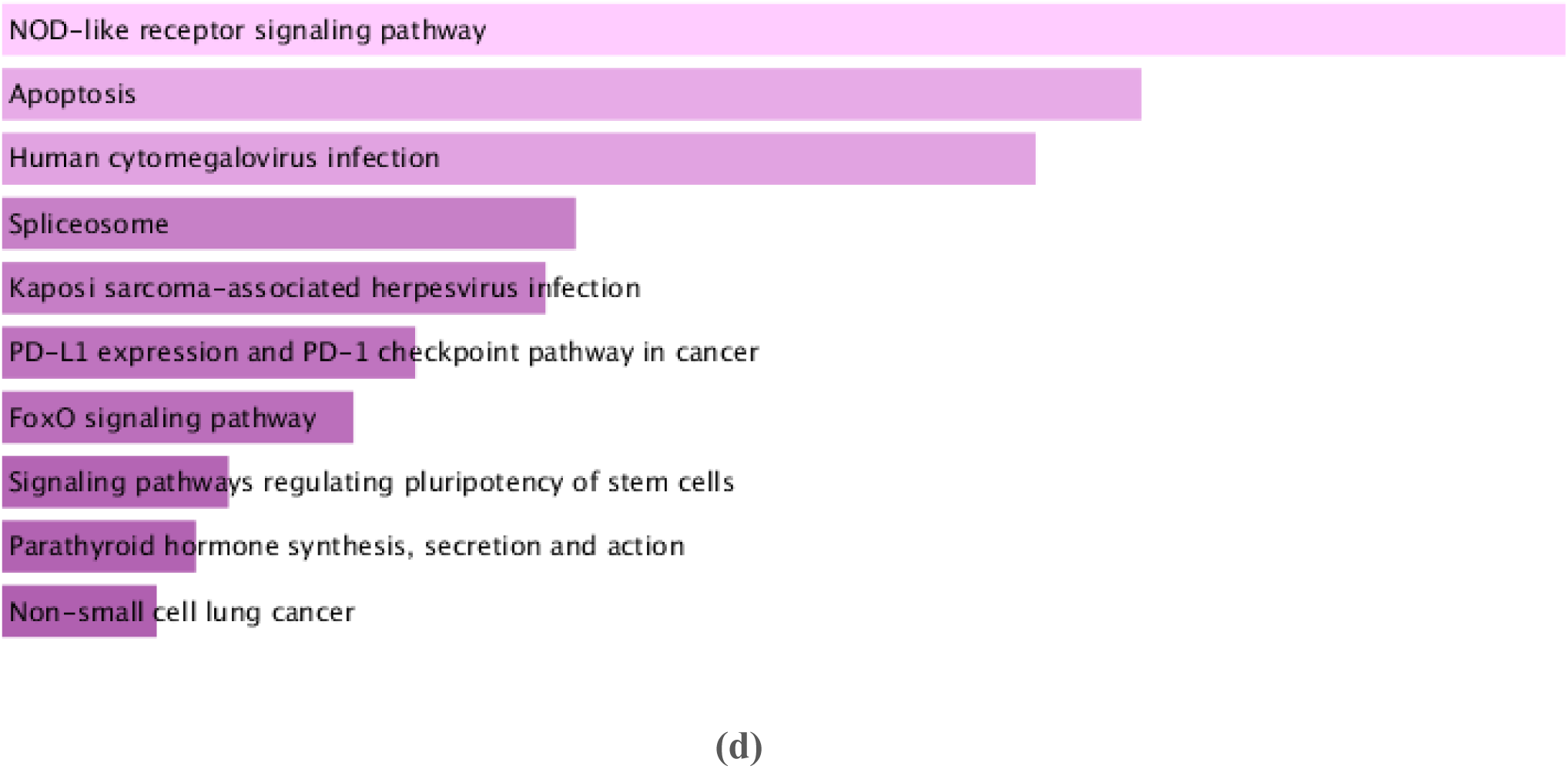
The enrichment analysis results in a graphical format. (a) The top ten biological processes enriched in the chosen set of DEGs. The x-axis represents the number of genes and the y-axis represents biological processes. (b) Top ten molecular functions that are enriched in these datasets. The x-axis represents the number of genes and the y-axis represents the molecular functions. (c) The top 10 cellular components are enriched in the DEGs of interest. The x-axis represents the number of genes and the y-axis represents the cellular components. (d) The DEGs of interest are enriched in the top 10 cellular components. The x-axis represents the number of genes and the y-axis represents the enriched KEGG pathways.

#### 3.5.2. Common GO molecular functions enriched

The molecular functions which are significantly enriched are RNA binding (GO:0003723) with a p-value of 0.000004154 followed by cytoskeleton nuclear membrane anchor activity (GO:0140444) with a p-value of 0.001746 and ubiquitin-protein transferase activity (GO:0004842) with a p-value of 0.002388. The molecular functions which are enriched with these are pre-mRNA binding, nuclear localization sequence binding, carbon-nitrogen ligase activity, with glutamine as amino-N-donor, pre-mRNA binding, Lys-63 specific deubiquitinase activity, protein kinase binding, and MAP-kinase. The graphical representation of the enriched molecular functions has been represented in Fig 8b.

#### 3.5.3. Common GO cellular components enriched

The cellular components which are enriched are the nuclear pathway (GO:0005634) with a p-value of 1.019e-10 followed by the pathways related to intracellular membrane-bound organelle (GO:00043231) with a p-value of 2.730e-8 and pathways associated with the condensed chromosome (GO:0000793) with a p-value of 0.0007446. The other cellular components include nuclear membrane, pericentric heterochromatin, prespliceosome, U2 type prespliceosomes, specific granule, phagocytic vesicle membrane, and U1 snRNP. Fig 8c illustrates the graphical display of enhanced cellular components.

#### 3.5.4. Common KEGG pathways that are enriched

The pathways named NOD-like receptor signaling pathway with a p-value of 0.0001612, followed by apoptotic pathway (p-value of 0.0006376), pathways involved in Human cytomegalovirus infection (p-value of 0.0008984), spliceosome, Kaposi’s sarcoma-associated herpesvirus infection, PD-L1 expression, and PD1 checkpoint pathway in cancer, FoxO signaling pathway, Signaling pathways regulating pluripotency of stem cells, Parathyroid hormone synthesis, secretion and action, and Non-small cell lung cancer. Figure 8d shows a graphical representation of enriched cellular components.

## 4. Discussion

The molecular pathways and genes of NDs are complex and in this study, we wanted to study the similarities in the expression of genes, the number of proteins that are common to these two diseases, miRNAs associated and also the pathways, biological processes, cellular component, and molecular functions associated with these two diseases.

The top genes that are found to be associated with both the datasets of these two diseases are PTPRC, CXCL8, RBM25. PTPRC encodes a protein called CD45, which is a protein tyrosine phosphatase that is known to regulate several pathways such as those associated with T-cell coactivation when it binds to the Dipeptidyl peptidase 4 (DPP4) a surface glycoprotein receptor (Ikushima et al., 2000). CD45 is a transmembrane protein with a single transmembrane domain. A point mutation in the PTPRC gene increases the risk for MS (Jacobsen et al., 2000). In another interesting study, it was seen that there was no connection between cytosine to guanine transition in the PTPRC gene and MS (Barcellos et al., 2001). CXCL8 is a gene that encodes Interleukin 8 (IL8) which according to a study was seen to elevate in HD patients (Björkqvist et al., 2008). In another study in which cerebrospinal fluid from MS patients was used as the study material, IL8 was elevated in MS patients compared to controls (Matejčíková et al., 2017). In a different study, lower levels of IL8 were seen in the CSF of Alzheimer’s disease patients, which is another critical ND (Hesse et al., 2016). RBM25 gene encodes a protein called RNA binding motif protein 25 which is a tumor suppressor protein. In a study, it was reported that knockdown of RBM25 results in the immortality of cells (by evading the apoptotic pathways) and also promotes proliferation in acute myeloid leukemia patients (Ge et al., 2019). In another study, it was found that RBM25 modulates the splicing of Bcl-x protein (Zhou et al., 2008). The role of RBM25 in MS and HD is yet to be established and thus this is a novel protein that should be studied to understand whether this affects these two NDs or not.

MicroRNAs are a key factor in neurodegenerative illnesses, and as more research is conducted, we are learning more about their function in the progression of these diseases (Goodall et al., 2013). The network that has been developed by us in this study showed that the genes that interact with maximum microRNAs and other genes are the MCL1 followed by the CXCL8 and NFKBIA (Fig 7). MCL1 is an antiapoptotic protein that belongs to the Bcl-2 family. In our current study, it was seen that MCL1 interacts with 17 other microRNAs, transcription factors, and genes. Higher expression of MCL1 in T cells was found in a study of MS patients, implying that dysregulation of the anti-apoptotic protein MCL1 may play a role in the disease’s progression (Mandel et al., 2012). CXCL8 is a gene that encodes IL8 which is associated with 16 other genes, transcription factors, and micro RNAs. NFKBIA encodes a protein that belongs to the NF-kappa-B inhibitor family. In a study, it was confirmed that MS patients having a mutation in the gene did not respond well to MIS416 treatment (McCombe et al., 2017). In this study, the NFKBIA interacts with ten other genes, transcription factors, and microRNAs. MicroRNAs such as hsa-mir-155-5p, hsa-mir-19a-3p, hsa-mir-93-5p, and hsa-mir-96-3p have maximum connectivity to other genes, transcription factors, and microRNAs in the network developed by us. In one study it was found out that microRNA by the name hsa-mir-93-5p has been elevated in the peripheral mononuclear blood samples of MS patients (Uwatoko et al., 2019). A microRNA hsa-mir-19b-3p which has a degree of 4 in the network created by us has been shown to upregulate in Parkinson’s disease patients (Uwatoko et al., 2019). The role of these upregulated microRNA in our current study needs more study for ascertaining their role in the development and progression of these NDs.

The 266 genes which are differentially expressed affect KEGG pathways that are associated with diabetic cardiomyopathy, neuroactive ligand-receptor interaction, Wnt signaling, and Hippo signaling (Deretzi et al., 2011). The DEGs showed that biological processes like the chemical synaptic transmission (GO:0007268), extracellular matrix organization (GO:0030198), inorganic cation transmembrane transport (GO:0098662) are linked to the development of these NDs (Centonze et al., 2010) (Smith-Dijak et al., 2019). Cytokine activity (GO:0005125), endopeptidase activity (GO:0004175), organic anion transmembrane transporter activity (GO:0008514), and G-protein coupled receptor binding activity (are the molecular functions that are affected by these two NDs according to our study. In several studies, it has been established now that elevated levels of specific cytokines such as IL6, IL8, TNFα, IFNγ are present in patients with these NDs (Florindo, 2014) (Rocha et al., 2016).

## Conclusion

In conclusion, the PPI network developed by us shows some proteins which were present in both the datasets of MS and HD were interconnected in the network. Significant connections were found with the DEGs and the microRNAs especially the hsa-mir-155-5p which has the highest degree. The enriched KEGG pathways showed that the Nod-like receptor signaling pathway is the most enriched pathway of all. Hence this study can work as a primer for other academic researches which would unravel more valuable information on both these NDs.

## List of abbreviations

ND: Neurodegenerative diseases
MS: Multiple sclerosis
HD: Huntington’s disease
DEGs: Differentially Expressed Genes

## Acknowledgments

None

